# *In silico* assessment of immune cross protection between BCoV and SARS-CoV-2

**DOI:** 10.1101/2024.01.25.577193

**Authors:** Lana Bazan Peters Querne, Fernanda Zettel Bastos, Mikaela dos Anjos Adur, Vitória Cavalheiro, Breno Castello Branco Beirão

## Abstract

**Background:** Humans have long shared infectious agents with cattle, and the bovine-derived human common cold OC-43 CoV is a not-so-distant example of cross-species viral spill over of coronaviruses. Human exposure to the Bovine Coronavirus (BCoV) is certainly common, as the virus is endemic in most high-density cattle-raising regions. Since BCoVs are phylogenetically close to SARS-CoV-2, it is possible that cross-protection against COVID-19 occurs in people exposed to BCoV.

**Methods:** This article shows an *in silico* investigation of human cross-protection to SARS-CoV-2 due to BCoV exposure. We determined HLA recognition and human B lymphocyte reactivity to BCoV epitopes using bioinformatics resources. A retrospective geoepidemiological analysis of COVID-19 was then performed to verify if BCoV/SARS-CoV-2 cross-protection could have occurred in the field. Brazil was used as a model for the epidemiological analysis of the impact of livestock density – as a proxy for human exposure to BCoV – on the prevalence of COVID-19 in people.

**Results:** As could be expected from their classification in the same *Betacoronavirus* genus, we show that several human B and T epitopes are shared between BCoV and SARS-CoV-2. This raised the possibility of cross-protection of people from exposure to the bovine coronavirus. Analysis of field data added partial support to the hypothesis of viral cross-immunity from human exposure to BCoV. There was a negative correlation between livestock geographical density and COVID-19. Whole-Brazil data showed areas in the country in which COVID-19 prevalence was disproportionally low (controlled by normalization by transport infrastructure). Areas with high cattle density had lower COVID-19 prevalence in these low-risk areas.

**Conclusions:** These data are hypothesis-raising indications that cross-protection is possibly being induced by human exposure to the Bovine Coronavirus.

## 1. Background

In December 2019 the Severe Acute Respiratory Syndrome Coronavirus 2 (SARS-CoV-2) was discovered in Wuhan, in the Chinese province of Hubei (1). SARS-CoV-2 can cause Coronavirus Disease 2019 (COVID-19) and led to a pandemic pneumonia outbreak, declared on March 11, 2020 (2). The symptoms of infected people resemble those of viral pneumonia, such as cough, fever and discomfort when breathing (3). In elderly patients and patients with comorbidities (e.g., diabetes, obesity, and asthma) the development of severe cases with dyspnoea and bilateral pulmonary infiltration is more common, increasing the number of hospitalizations and deaths in this population (4).

Coronaviruses are single-stranded RNA viruses belonging to the *Coronaviridae* family, which infects several animal hosts. Within this range of hosts, coronaviruses cause respiratory, gastrointestinal, and neurological diseases. The four genera that compose this family are: *Alphacoronavirus*, *Betacoronavirus*, *Gammacoronavirus* and *Deltacoronavirus* (5,6). Among the Beta-coronaviruses are SARS-CoV-2 and Bovine Coronavirus (BCoV). The latter is responsible for livestock losses, causing diarrhoea in new-born calves and respiratory infections in calves and confined cattle (6,7). The genome of both viruses encodes similar structural proteins: envelope protein (E), membrane protein (M), nucleocapsid protein (N) and spike protein (S); BCoV expresses a hemagglutinin-esterase not present in SARS-CoV-2 (8); viruses also express homologous non-structural proteins (NSP) and open reading frame polyproteins (ORF) (6,9).

Cross-reactivity between coronaviruses is known to occur to some extent and might impact on the severity and spread of diseases (10). BCoV and SARS-CoV-2 are aggregated within the same viral genus which illustrates the high structural similarity between them – this is crucial for immune cross-reactivity (11,12). Importantly, there is a history of BCoV spill over to other species, including humans, which seems to have generated at least one of the current human coronaviruses that cause the common cold (8). It is possible that subclinical human infections with BCoV occur routinely, and there is even evidence of BCoV causing clinical signs in susceptible people (13,14).

Here, we performed an *in silico* analysis of the correlations between bovine and human coronaviruses. We conducted an immunological assessment of the epitopes of BCoV which may induce protective immune responses in humans against SARS-CoV-2. We searched for peptides originated from BCoV proteins M, N, S and ORF that potentially could induce T and B cell responses in people and that show high identity with SARS-CoV-2. We then used an epidemiological analysis to test the hypothesis that exposure to BCoV induces cross-protection against COVID-19 (cattle density was used as a proxy for BCoV exposure). The results presented here are an indication that BCoV may confer human cross-protection against SARS-CoV-2.

## 2. Methods

### 2.1 Peptide setup for immunological assessment

Proteome sequences of Bovine Coronavirus were obtained from the NCBI database and focused on four proteins (Table 1): spike protein (S), membrane protein (M), nucleocapsid protein (N) and replicase polyprotein (Orf1ab). The entire protein sequences were organized in 15-mer peptides that overlapped by 10 amino acids, using a python code (15,16).

**Table 1.** NCBI accession numbers of Bovine Coronavirus and SARS-CoV-2 protein sequences used in the present study.

|  | **Bovine coronavirus** | **SARS-CoV-2** |
| --- | --- | --- |
| Spike protein | NP_150077.1 | YP_009724390.1 |
| Membrane protein | NP_150082.1 | YP_009724393.1 |
| Nucleocapsid protein | NP_150084.1 | QQD86936.1 |
| Orf1ab | NP_150073.3 | BCT04066.1 |

Orf1ab = replicase polyprotein.

### 2.2 Prediction of T cell reactivity

T cell reactivity of bovine coronavirus peptides was assessed by predicting their binding to human leukocyte antigen class II (HLA II) molecules using IEDB MHC II binding predictions tool (http://tools.iedb.org/mhcii/). Peptide binding was predicted to all HLA class II molecules. A 20% percentile rank cut-off was chosen as a universal prediction threshold (16).

### 2.3 Prediction of B cell reactivity

B cell reactivity of bovine coronavirus peptides was assessed using IEDB Bepipred Linear Epitope Prediction 2.0 (http://tools.iedb.org/bcell/). The residues with scores above the threshold (0.5) and with 5 amino acids or more were predicted to be part of an epitope (17,18).

### 2.4 Similarity of BCoV peptides in relation to SARS-CoV-2 proteins

All BCoV peptides that were above the thresholds in the analyses of T- and B cells were assessed for their similarity to the corresponding proteins of SARS-CoV-2 (Table 1) using the Multiple Sequence Alignment (Clustal Omega, https://www.ebi.ac.uk/Tools/msa/clustalo/). Sequences with an identity greater than or equal to 80% were selected as peptide matches (19).

### 2.5 Epidemiology of COVID-19 and association with BCoV

Spatial correlation between cattle and COVID-19 was assessed using data from Brazil. The country has large and well-defined areas with high cattle density. Also, within-state analyses allow for controlled comparison of COVID-19 risk factors, as the most important public policies that alter COVID-19 risks are more homogeneously distributed in a state level (20).

COVID-19 epidemiology was assessed from publicly available data (21). For a within-state analysis, the slope of increase of cases/100,000 people for each city in the Brazilian State of Mato Grosso do Sul (MS) was used (between January, 2020 and September, 2021) (21,22). The slope of COVID-19 cases was compared to the number of cattle/100,000 people for each municipality in the state (23).

As a control, the distance from each municipality to the major city in the subregion of the state was compared to the slope of COVID-19 cases (24). General efficiency of public spending (not directly correlated with COVID-19) was also used as a control in a correlation analysis with COVID-19 prevalence. Data from the literature on public investment were used. Spending rigor was scored from 1-4, with four being the best-quality public use of resources (25). The correlation of the data with COVID-19 prevalence was assessed with run’s test in a linear correlation.

Whole-country data from Brazil was assessed using QGIS 3.24.1 Tisler. COVID-19 data from every Brazilian municipality and the map of Brazilian roads were obtained from the Instituto Brasileiro de Geografia e Estatística (26). Cattle population localization and density was from a previously published dataset (27).

COVID-19 prevalence rates, road density and cattle populations were compared by pixel intensity of the respective rasterized layers using ‘Point Sampling Tool Plugin’ for QGIS (version 0.5.3, by Borys Jurgiel). A grid of dots was layered on top of the maps of interest for analysis using the plugin. The grid was positioned to cover the entirety of Brazilian territory south of the Equator, where cattle-raising regions are located. COVID-19 prevalence was corrected in relation to road density in the respective region. For this, every COVID-19 dot from the analysis grid was divided by the sum of the 9 surrounding road ‘intensity’ dots.

Raw data used for epidemiological analysis is provided as a supplementary material.

GraphPad Prism 8 (GraphPad Software, Inc., USA) was used for graphing and for statistical analysis. All the data used for this analysis is available as supplementary material.

## 3. Results

### 3.1 Peptide setup for immunological assessment

A total of 136, 23, 45 and 709 15-mer peptides that overlapped by 10 amino acids were obtained for proteins S, M, N and ORF1ab respectively.

### 3.2 Prediction of T cell reactivity

From the results obtained by the IEDB MHC II binding prediction tool, 106 peptides from protein S, 20 peptides from protein M, 24 peptides from protein N and 566 peptides from ORF1ab protein were above the selection threshold. All peptides obtained in this analysis are available as supplementary material.

### 3.3 Prediction of B cell reactivity

From the results obtained by the IEDB Bepipred Linear Epitope Prediction 2.0, 70 peptides from protein S, 9 peptides from protein M, 38 peptides from protein N and 386 peptides from ORF1ab protein had scores above the threshold. All peptides obtained in this analysis are available as supplementary material.

### 3.4 Similarity of BCoV peptides in relation to SARS-CoV-2 proteins

Among the peptides that showed good results for putative human T or B cell interactions, only 2 peptides from protein S, 1 peptide from protein M, and 2 peptides from protein N showed at least 80% similarity with SARS-CoV-2 (Table 2). No peptide sequence from these three proteins was found to be above the cut-off values for both T cells and B cells.

**Table 2.**
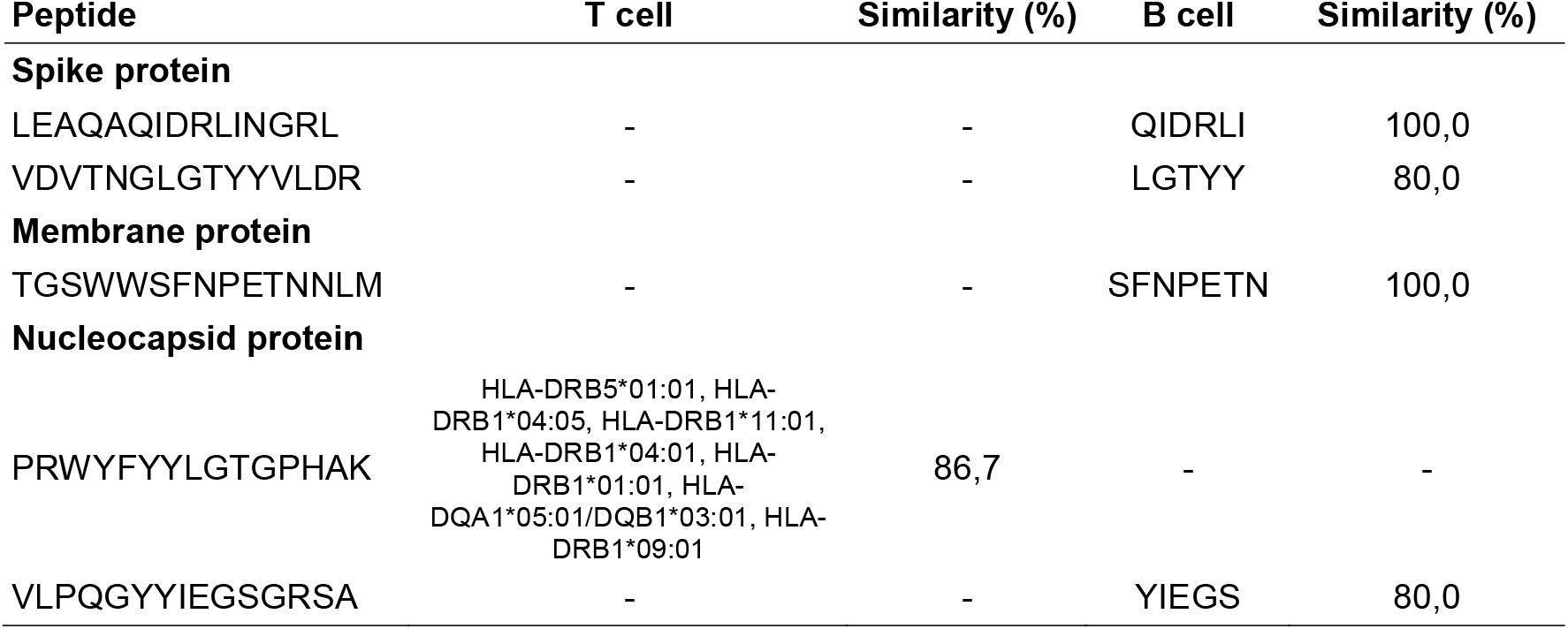
Peptides from BCoV spike, membrane and nucleocapsid proteins that were likely to induce human T- and B cell responses and that showed at least 80% similarity with SARS-CoV-2.

Regarding the ORF1ab protein,107 peptides were above the threshold for potential T- or B cell epitopes. In this case, 28 peptides were found to be above the cut-off for both T cells and B cells (Table 3).

**Table 3.**
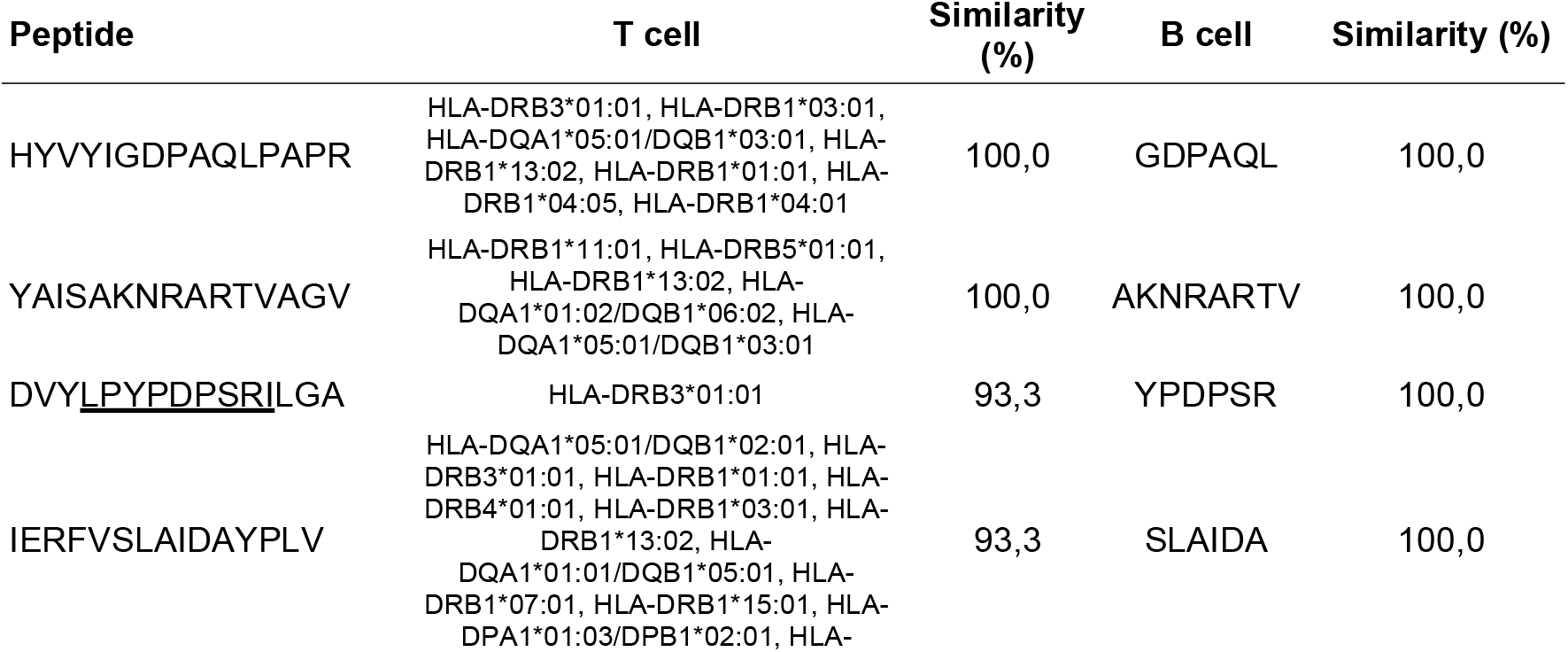
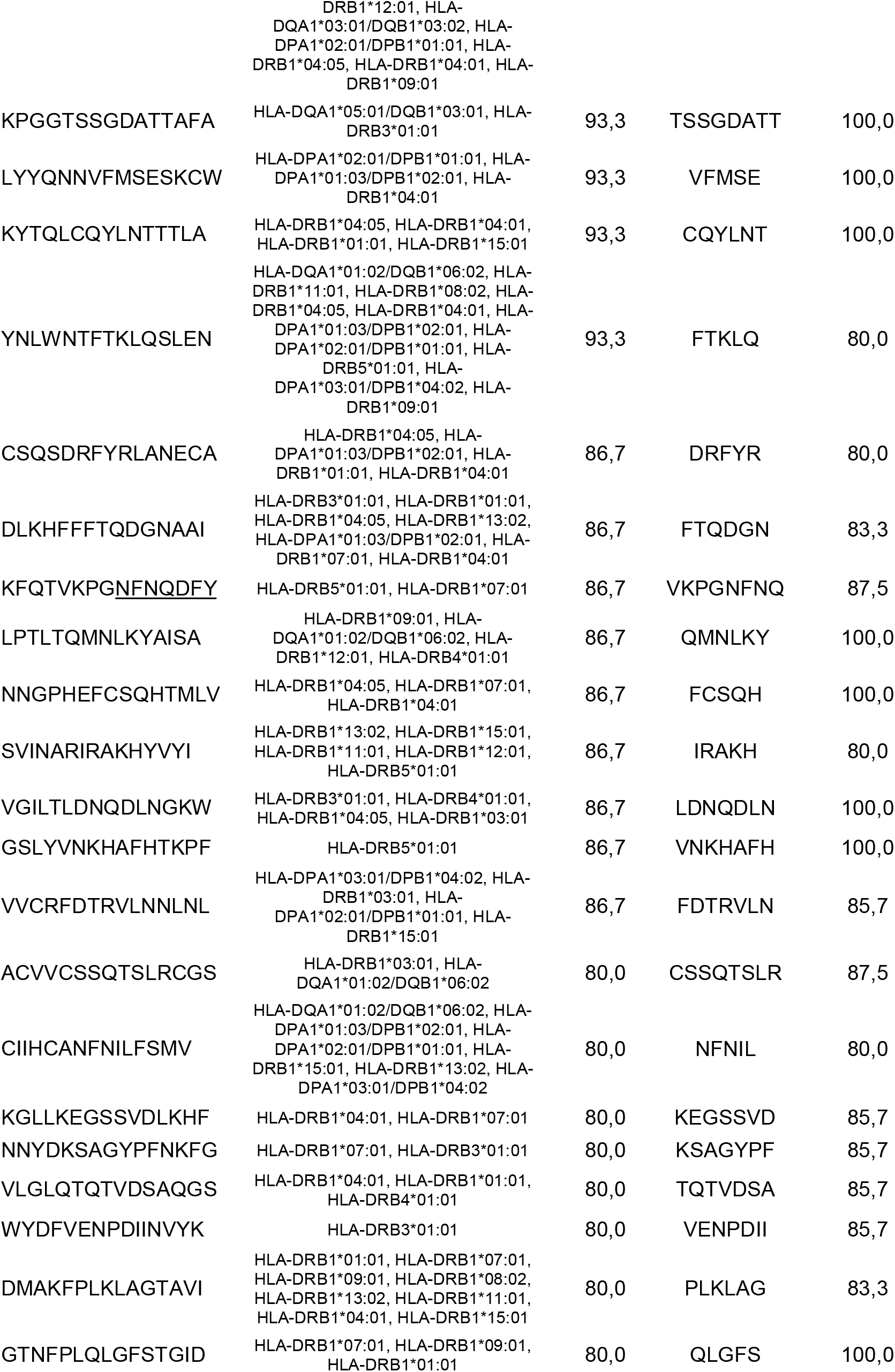
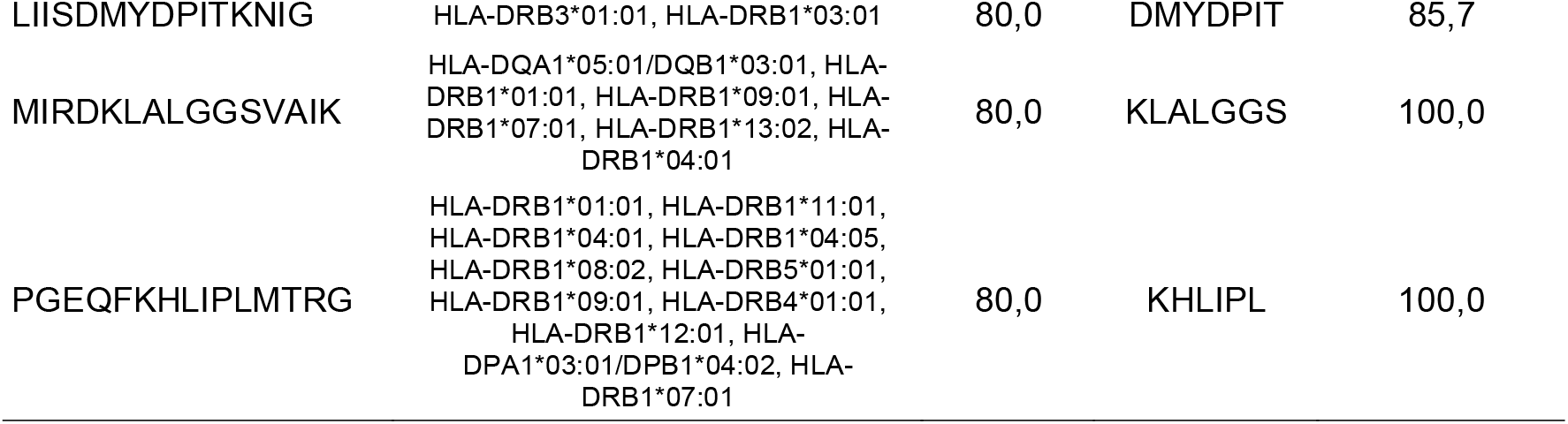
Peptides from replicase polyprotein (ORF1ab) that were likely to induce human T- and B cell responses and that showed at least 80% similarity with SARS-CoV-2. The underlined BCoV peptide sequence has been reported to induce protective anti-SARS-CoV-2 T cell responses (28).

### 3.5 Epidemiology of COVID-19 and association with BCoV

To test the hypothesis that BCoV exposure may lead to COVID-19 cross-reactive immunity, we performed an epidemiological assessment of the correlation between these factors. We analysed the correlation of COVID-19 prevalence to the density of cattle in the Brazilian state of MS. This was performed as an initial investigation into the epidemiological association between human exposure to the Bovine Coronavirus (BCoV) and altered pandemic spread. Cattle density was used as a proxy for BCoV exposure.

Cattle density (cattle/100,000 people) negatively correlated with the slope of COVID-19 case increase in MS. In opposition, confounding factors in this epidemiological analysis showed no association with the slope of COVID-19 cases in the state (assessed factors were distance of each municipality to the main regional hub city and quality of public spending) (Fig. 1).

**Figure 1.**
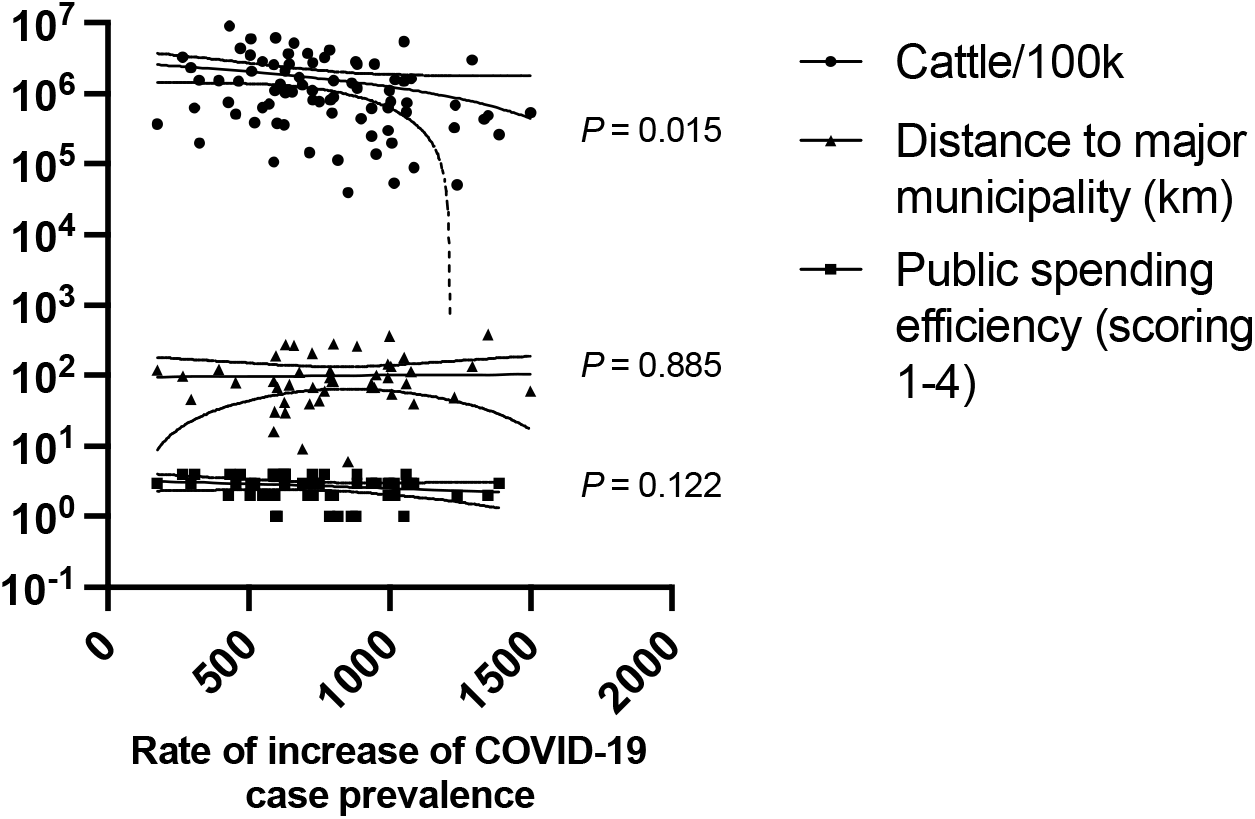
Linear regression between cattle density and the slope of cumulative COVID-19 case increase in the Brazilian State of Mato Grosso do Sul (MS). Data between Jan/20 and Sep/21 were used. Cattle density was calculated as the number of cattle/100,000 people in the municipality. The distance of the municipality to the major hub city was used to control for lower people connectivity of cattle-raising areas. Public spending efficiency was used to control for possible slower responses to the COVID-19 pandemic from cattle-raising municipalities. Analysis by run’s test in a linear regression. *P-*values are shown next to each regression. The dotted lines around the linear regression trend indicate the 99% CI.

In a second proof-of-concept epidemiological analysis, we determined the statistical correlation between a) COVID-19 prevalence throughout the country of Brazil; b) the density of cattle populations in the respective areas. As a normalizer for the data, human circulation was determined by assessing road density in the area.

Brazilian municipalities were classified as either a) having less COVID-19 cases than expected by the surrounding road infrastructure or, b) having more COVID-19 cases than expected by the surrounding road infrastructure. In the former low-risk cohort, bovine density was negatively correlated to COVID-19 prevalence (Fig. 2).

**Figure 2.**
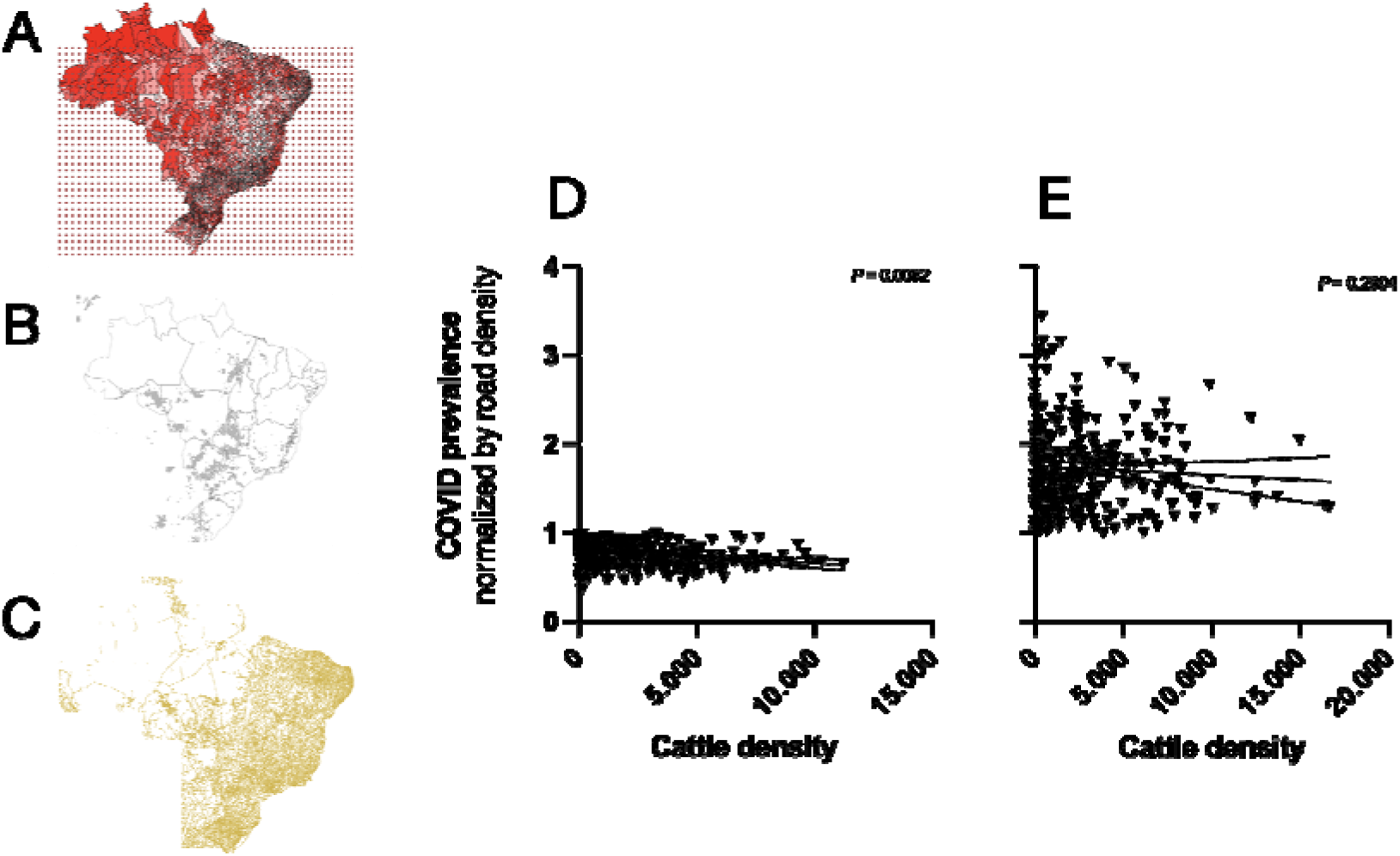
Cattle density was negatively associated to COVID-19 prevalence in Brazil. COVID-19 prevalence (A, in red) and cattle density (B, gray shades) were assessed using a geographic information system. Road density (C) was also assessed as a control for the relevance of human movement in COVID-19 prevalence. Although not shown, the dotted grid applied in (A) was also in (B) and (C) for the respective measurements. The effect of cattle density was assessed separately in areas with low (D) and high risk (E) (as expected from road density). The *P* value indicates departure from linearity for the correlation lines by run’s test. The dashed line indicates 95% CI.

## 4. Discussion

Bovine coronaviruses (BCoV) are members of the *Betacoronavirus* genus along with SARS-CoV-2, denoting their structural similarities. Further, within the *Betacoronavirus*, BCoV is among the most similar viruses to SARS-CoV-2 (29,30). Indeed, cattle can be experimentally infected with SARS-CoV-2 (31–33) and bovine coronaviruses have spilled over to humans before - current strains of BCoV can be cultured in human rectal adenocarcinoma cells, demonstrating that cross-species infection is still a risk, if not a common event already (6,34,35). Other works have already discussed the immunological impacts that coronaviruses of domestic animals could have on humans. In Brazil, the use of the *Deltacoronavirus* Avian Infectious Bronchitis is being clinically tested for COVID-19 vaccination, for instance [11, 24].

The hypothesis raised here is that BCoV exposure influences human immune responses to COVID-19. We started our evaluation of the cross-protection between BCoV and SARS-CoV by assessing *in silico* if BCoV epitopes could be recognized by human B and T lymphocytes. Here, we report several BCoV epitopes which are likely to be important in the immune response against COVID-19. This analysis is valuable in confirming that infectious exposure to the bovine coronavirus can theoretically induce SARS-CoV-2 cross-reactive immune responses – although it must be made clear that human infectivity of BCoV cannot be confirmed with the present analysis.

Since BCoV shares epitopes with SARS-CoV-2, it is possible that COVID-19 epidemiology was shaped by human exposure to BCoV, much as smallpox was naturally curtailed by the exposure to cowpox, for instance (36). BCoV naturally and widely occurs in densely populated bovine herds (14) and there is evidence of human transmission (37). In this scenario, BCoV exposure would be one among other interacting factors in COVID-19 spread, such as income and social vulnerability levels (38). The results from the Brazilian state of MS and the wider analysis of the country were supportive of the hypothesis that human exposure to cattle had an impact on the epidemiology of COVID-19.

The state of MS was chosen as a proof-of-concept case study, as it is a large beef productor with no megacities, which can “distort” the local epidemiological status due to their large influence on the statistics and due to their disproportionate worldwide connections in relation to other towns (39,40). For Brazil, within-state infrastructure, scholarity, income and animal production conditions are more homogeneous than in inter-state comparisons (41,42), thus explaining the choice of a state for the preliminary epidemiological analysis.

COVID-19 data from MS was compared against general efficiency of public spending – an important factor in the spread and control of the pandemic in Brazil (43) – and against distance to major city hubs. Municipalities with more cattle are expected to be further away from regional hubs, since large land areas are needed for extensive bovine farming. Therefore, any association between COVID-19 cases with cattle density could possibly be due to lower connectivity of the municipality, which is a major cause of spatial proliferation of the disease (44). These data were freely available and were therefore used for the analysis of the state of MS. Within MS, cattle density was negatively correlated to COVID-19 cases, being more significant in explaining pandemic expansion than common biases, public spending efficiency and distance to major cities.

Whole-country Brazilian COVID-19 data demonstrated an interesting pattern in which some municipalities were more “benefitted” from exposure to cattle. COVID-19 prevalence was corrected for road density, creating an index of cities that had higher or lower COVID-19 prevalence than theoretically expected based on road density (an inference of populational movements, which highly alter infectious disease spread (45,46)). “Lower-than-expected” COVID-19 rates could indicate a myriad of factors, such as better health systems or stricter municipal COVID-19 control laws. In this case, the results would indicate that BCoV exposure only benefited human populations that had low risks for COVID-19. High SARS-CoV-2 exposure may have overcome any benefits conferred by previous contact with BCoV. Regional road density may also not have appropriately normalized COVID-19 prevalence, as it is a single and limited control (47).

It must be stressed that it was not the goal of this study to prove the association of BCoV with COVID-19 using epidemiological data. Our analysis is exceedingly restricted for this purpose. Nevertheless, these results are an indication of immune cross-reactivity and potential protection from COVID-19 from exposure to BCoV. These data prompt further experimental analyses of the effect of BCoV in people.

## 5. Conclusion

SARS-CoV-2 and BCoV share several common epitopes, which may confer cross-immunity. The relevance of this for the development of the pandemic is yet not known and should be proven with controlled trials of human responses to the bovine virus. Nevertheless, our results for the correlation between COVID-19 prevalence and cattle density are an indication of the role of human exposure to BCoV with regards to the development of the pandemic.

## List of abbreviations

BCoV: Bovine Coronavirus
MS: Mato Grosso do Sul [State of Brazil]

## Ethical approval and consent to participate

Not required.

## Consent for publication

Not applicable.

## Availability of data and materials

All data generated or analysed during this study are included in this published article [and its supplementary information files].

## Author contributions

LBPQ and FZB performed the molecular analyses with viral genomic data. VC, MAA and BCBB performed the epidemiological analyses. LBPQ, FZB and BCBB wrote the article. BCBB provided funding for the research.

## Acknowledgments

Not applicable.

## References

1. Walls AC, Park YJ, Tortorici MA, Wall A, McGuire AT, Veesler D. Structure, Function, and Antigenicity of the SARS-CoV-2 Spike Glycoprotein. Cell. 2020 Apr 16;181(2):281–292.e6.

2. WHO Coronavirus (COVID-19) Dashboard | WHO Coronavirus (COVID-19) Dashboard With Vaccination Data [Internet]. [cited 2023 Jun 25]. Available from: https://covid19.who.int/

3. Chen Y, Li L. SARS-CoV-2: virus dynamics and host response. Lancet Infect Dis [Internet]. 2020 May 1 [cited 2023 Jun 25];20(5):515–6. Available from: https://pubmed.ncbi.nlm.nih.gov/32213336/

4. Hu B, Guo H, Zhou P, Shi ZL. Characteristics of SARS-CoV-2 and COVID-19. Nature Reviews Microbiology 2020 19:3 [Internet]. 2020 Oct 6 [cited 2023 Jun 25];19(3):141–54. Available from: https://www.nature.com/articles/s41579-020-00459-7

5. Wu D, Wu T, Liu Q, Yang Z. The SARS-CoV-2 outbreak: What we know. International Journal of Infectious Diseases [Internet]. 2020 May 1 [cited 2023 Jun 25];94:44. Available from: /pmc/articles/PMC7102543/

6. Vlasova AN, Saif LJ. Bovine Coronavirus and the Associated Diseases. Front Vet Sci. 2021 Mar 31;8.

7. Takiuchi E, Alfieri AF, Alfieri AA. Molecular analysis of the bovine coronavirus S1 gene by direct sequencing of diarrheic fecal specimens. Braz J Med Biol Res [Internet]. 2008 [cited 2023 Jun 25];41(4):277–82. Available from: https://pubmed.ncbi.nlm.nih.gov/18392449/

8. Lang Y, Li W, Li Z, Koerhuis D, Van Den Burg ACS, Rozemuller E, et al. Coronavirus hemagglutinin-esterase and spike proteins coevolve for functional balance and optimal virion avidity. [cited 2023 Jun 25]; Available from: https://www.pnas.org

9. Kung YA, Lee KM, Chiang HJ, Huang SY, Wu CJ, Shih SR. Molecular Virology of SARS-CoV-2 and Related Coronaviruses. Microbiology and Molecular Biology Reviews. 2022 Jun 15;86(2).

10. Ellis J, Sniatynski M, Rapin N, Lacoste S, Erickson N, Haines D. SARS coronavirus 2-reactive antibodies in bovine colostrum. Can Vet J [Internet]. 2023 [cited 2023 Jun 24];337–43. Available from: https://www.ncbi.nlm.nih.gov/pmc/articles/PMC10031788

11. Lee CH, Pinho MP, Buckley PR, Woodhouse IB, Ogg G, Simmons A, et al. Potential CD8+ T cell cross-reactivity against SARS-CoV-2 conferred by other coronavirus strains. Front Immunol. 2020;11:579480.

12. What happened to the Covid vaccines Brazil promised to develop? [Internet]. [cited 2023 Jul 6]. Available from: https://brazilian.report/society/2023/06/09/covid-vaccines-promised-develop/

13. Virant MJ, Černe D, Petrovec M, Paller T, Toplak I. Genetic characterisation and comparison of three human coronaviruses (Hku1, oc43, 229e) from patients and bovine coronavirus (bcov) from cattle with respiratory disease in Slovenia. Viruses [Internet]. 2021 Apr 1 [cited 2023 Jun 25];13(4). Available from: /pmc/articles/PMC8071153/

14. Zhu Q, Li B, Sun D. Advances in Bovine Coronavirus Epidemiology. Viruses [Internet]. 2022 May 1 [cited 2023 Jun 25];14(5):1109. Available from: https://www.mdpi.com/1999-4915/14/5/1109/htm

15. Sanchez-Trincado JL, Gomez-Perosanz M, Reche PA. Fundamentals and Methods for T- and B-Cell Epitope Prediction. J Immunol Res [Internet]. 2017 [cited 2023 Jul 6];2017. Available from: https://pubmed.ncbi.nlm.nih.gov/29445754/

16. Paul S, Sidney J, Peters B, Sette A. Development and validation of a broad scheme for prediction of HLA class II restricted T cell epitopes. In: Proceedings of the 5th ACM Conference on Bioinformatics, Computational Biology, and Health Informatics. 2014. p. 733–8.

17. Gasteiger E, Hoogland C, Gattiker A, Duvaud S, Wilkins MR, Appel RD, et al. Protein identification and analysis tools on the ExPASy server. Springer; 2005.

18. Jespersen MC, Peters B, Nielsen M, Marcatili P. BepiPred-2.0: improving sequence-based B-cell epitope prediction using conformational epitopes. Nucleic Acids Res [Internet]. 2017 Jul 3 [cited 2023 Jul 6];45(W1):W24–9. Available from: 10.1093/nar/gkx346

19. Reche PA. Potential cross-reactive immunity to SARS-CoV-2 from common human pathogens and vaccines. Front Immunol. 2020;2694.

20. Informações básicas municipais [Internet]. [cited 2023 Jun 25]. Available from: http://tabnet.fiocruz.br/dhx.exe?observatorio/fat_indicadores.def

21. Covid19 por Município - Brasil.IO [Internet]. [cited 2023 Jul 6]. Available from: https://brasil.io/covid19/

22. Islam ARMT, Hasanuzzaman M, Shammi M, Salam R, Bodrud-Doza M, Rahman MM, et al. Are meteorological factors enhancing COVID-19 transmission in Bangladesh? Novel findings from a compound Poisson generalized linear modeling approach. Environmental Science and Pollution Research. 2021;28:11245–58.

23. Pesquisa da Pecuária Municipal | IBGE [Internet]. [cited 2023 Jul 6]. Available from: https://www.ibge.gov.br/estatisticas/economicas/agricultura-e-pecuaria/9107-producao-da-pecuaria-municipal.html

24. Estudo da Dimensão Territorial do Estado de MS: Regiões de Planejamento – SEMADESC [Internet]. [cited 2023 Jul 6]. Available from: https://www.semadesc.ms.gov.br/estudo-da-dimensao-territorial-do-estado-de-ms-regioes-de-planejamento/

25. Dorsa ACC, Taveira JC, Pereira MS, Santos FK, Costa RB. Eficiência dos municípios de Mato Grosso do Sul: uma abordagem baseada em fronteira determinística. Interações (Campo Grande). 2020;21:663–80.

26. Portal de mapas do IBGE [Internet]. [cited 2022 Apr 15]. Available from: https://portaldemapas.ibge.gov.br/portal.php#homepage

27. Gilbert M, Nicolas G, Cinardi G, Van Boeckel TP, Vanwambeke SO, Wint GRW, et al. Global distribution data for cattle, buffaloes, horses, sheep, goats, pigs, chickens and ducks in 2010. Sci Data. 2018;5.

28. Swaminathan S, Lineburg KE, Ambalathingal GR, Crooks P, Grant EJ, Mohan S V., et al. Limited Recognition of Highly Conserved Regions of SARS-CoV-2. Microbiol Spectr [Internet]. 2022 Feb 23 [cited 2023 Jun 25];10(1). Available from: https://journals.asm.org/doi/10.1128/spectrum.02780-21

29. Zhou H, Ji J, Chen X, Bi Y, Li J, Wang Q, et al. Identification of novel bat coronaviruses sheds light on the evolutionary origins of SARS-CoV-2 and related viruses. Cell [Internet]. 2021 Aug 19 [cited 2023 Jul 6];184(17):4380–4391.e14. Available from: https://pubmed.ncbi.nlm.nih.gov/34147139/

30. Tilocca B, Soggiu A, Musella V, Britti D, Sanguinetti M, Urbani A, et al. Molecular basis of COVID-19 relationships in different species: a one health perspective. Microbes Infect. 2020 May 1;22(4–5):218–20.

31. Ulrich L, Wernike K, Hoffmann D, Mettenleiter TC, Beer M. Experimental Infection of Cattle with SARS-CoV-2. Emerg Infect Dis [Internet]. 2020 Dec 1 [cited 2023 Jul 6];26(12):2979–81. Available from: https://pubmed.ncbi.nlm.nih.gov/33034284/

32. Fusco G, Cardillo L, Levante M, Brandi S, Picazio G, Napoletano M, et al. First serological evidence of SARS-CoV-2 natural infection in small ruminants: Brief report. Vet Res Commun. 2023;

33. Bosco-Lauth AM, Walker A, Guilbert L, Porter S, Hartwig A, McVicker E, et al. Susceptibility of livestock to SARS-CoV-2 infection. Emerg Microbes Infect. 2021;10(1):2199–201.

34. Brüssow H, Brüssow L. Clinical evidence that the pandemic from 1889 to 1891 commonly called the Russian flu might have been an earlier coronavirus pandemic. Microb Biotechnol [Internet]. 2021 Sep 1 [cited 2023 Jul 6];14(5):1860–70. Available from: https://pubmed.ncbi.nlm.nih.gov/34254725/

35. Stipp DT. Detecção do coronavírus bovino em episódios de diarréia neonatal em rebanhos bovinos brasileiros. 2007;

36. Riedel S. Edward Jenner and the history of smallpox and vaccination. Proc (Bayl Univ Med Cent) [Internet]. 2005 Jan 1 [cited 2023 Jul 6];18(1):21. Available from: /pmc/articles/PMC1200696/

37. Zhu Q, Li B, Sun D. Advances in Bovine Coronavirus Epidemiology. Viruses [Internet]. 2022 May 1 [cited 2023 Jun 25];14(5). Available from: https://www.ncbi.nlm.nih.gov/pmc/articles/PMC9147158

38. Cestari VRF, Florêncio RS, Sousa GJB, Garces TS, Maranhão TA, Castro RR, et al. Social vulnerability and COVID-19 incidence in a Brazilian metropolis. Cien Saude Colet [Internet]. 2021 Mar 1 [cited 2023 Jul 6];26(3):1023–33. Available from: https://pubmed.ncbi.nlm.nih.gov/33729356/

39. Ren H, Zhao L, Zhang A, Song L, Liao Y, Lu W, et al. Early forecasting of the potential risk zones of COVID-19 in China’s megacities. Science of the Total Environment. 2020 Aug 10;729.

40. Urban densities and the Covid-19 pandemic: Upending the sustainability myth of global megacities | ORF [Internet]. [cited 2023 Jul 6]. Available from: https://www.orfonline.org/research/urban-densities-and-the-covid-19-pandemic-upending-the-sustainability-myth-of-global-megacities-65606/

41. Lebioda L, Cabral GO, Tezza R. A Homogeneidade da Inclusão Digital no Brasil: Sonho ou Realidade? Revista Informação na Sociedade Contemporânea [Internet]. 2019 Dec 30 [cited 2023 Jul 6];3(1):1–18. Available from: https://periodicos.ufrn.br/informacao/article/view/19118

42. Krawczyk NR, Vieira VL. Homogeneidade e heterogeneidade nos sistemas educacionais: Argentina, Brasil, Chile e México. Cadernos de Pesquisa [Internet]. 2006 Sep [cited 2023 Jul 6];36(129):673–704. Available from: https://www.scielo.br/j/cp/a/VJN8jzhCtnBpkkyMPs4qD6y/?lang=pt

43. Szylovec A, Umbelino-Walker I, Cain BN, Ng HT, Flahault A, Rozanova L. Brazil’s Actions and Reactions in the Fight Against COVID-19 from January to March 2020. Int J Environ Res Public Health [Internet]. 2021 Jan 2 [cited 2023 Jul 6];18(2):1–16. Available from: https://pubmed.ncbi.nlm.nih.gov/33440812/

44. Jo Y, Hong A, Sung H. Density or Connectivity: What Are the Main Causes of the Spatial Proliferation of COVID-19 in Korea? Int J Environ Res Public Health [Internet]. 2021 May 2 [cited 2023 Jul 6];18(10). Available from: https://pubmed.ncbi.nlm.nih.gov/34065031/

45. Khavarian-Garmsir AR, Sharifi A, Moradpour N. Are high-density districts more vulnerable to the COVID-19 pandemic? Sustain Cities Soc. 2021 Jul 1;70:102911.

46. Guan C, Tan J, Hall B, Liu C, Li Y, Cai Z. The Effect of the Built Environment on the COVID-19 Pandemic at the Initial Stage: A County-Level Study of the USA. Sustainability (Switzerland) [Internet]. 2022 Mar 1 [cited 2023 Jun 25];14(6):3417. Available from: https://www.mdpi.com/2071-1050/14/6/3417/htm

47. Coura-Vital W, Cardoso DT, Ker FT de O, Magalhães FDC, Bezerra JMT, Viegas AM, et al. Spatiotemporal dynamics and risk estimates of COVID-19 epidemic in Minas Gerais State: analysis of an expanding process. Rev Inst Med Trop Sao Paulo [Internet]. 2021 Mar 24 [cited 2022 Apr 15];63. Available from: http://www.scielo.br/j/rimtsp/a/yfqPmQ5SyDCscbmy7gbhFsD/?lang=en

