## Supplementary figures and images for "*In silico* assessment of immune cross protection between BCoV and SARS-CoV-2"

### Brazil COVID.tiff

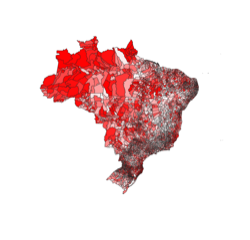

### Estradas Brasil em raster.tif

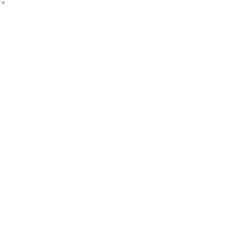
